# Large-scale, dynamin-like motions of the human guanylate binding protein 1 revealed by multi-resolution simulations

**DOI:** 10.1101/676981

**Authors:** Bogdan Barz, Jennifer Loschwitz, Birgit Strodel

## Abstract

Guanylate binding proteins (GBPs) belong to the dynamin-related superfamily and exhibit various functions in the fight against infections. The functions of the human guanylate binding protein 1 (hGBP1) are tightly coupled to GTP hydrolysis and dimerization. Despite known crystal structures of the hGBP1 monomer and GTPase domain dimer, little is known about the dynamics of hGBP1. To gain a mechanistic understanding of hGBP1, we performed sub-millisecond multi-resolution molecular dynamics simulations of both the hGBP1 monomer and dimer. We found that hGBP1 is a highly flexible protein that undergoes a hinge motion similar to the movements observed for other dynamin-like proteins. Another large-scale motion was observed for the C-terminal helix *α*13, providing a molecular view for the *α*13–*α*13 distances previously reported for the hGBP1 dimer. Most of the loops of the GTPase domain were found to be flexible, disclosing why GTP binding is needed for hGBP1 dimerization to occur.

**Author summary:** Gunaylate binding proteins are key fighters against microbial and viral pathogens. In the human body there are seven types of such proteins, among which is the guanylate binding protein 1 (hGBP1). This protein is able to perform its function only once it is activated by binding and converting guanosinetriphosphat (GTP) to guanosinediphosphat and guanosinemonophosphat via hydrolysis. In concert with the conversion of GTP the dimerization of hGBP1 occurs, which can further interact with the lipid membrane of the pathogen and disrupt it. While the crystal structure of the protein is known, the activation and dimerization steps are not well understood at molecular level as studying them experimentally is difficult. An alternative approach is given by molecular simulations, allowing us to elucidate the protein dynamics closely connected to these steps. From our simulations applied to both the hGBP1 monomer and dimer we identified large-scale motions taking place in hGBP1 that had not been reported before. We discuss the relevance of these motions in terms of their biological function, such as possible membrane damage caused by one of the motions or locking the protein in the dimer state.

## Introduction

Guanosine triphosphate (GTP) binding proteins play essential roles in many cellular processes responsible for the maintenance and regulation of biological functions. Among these proteins are the guanylate binding proteins (GBPs), which belong to the dynamin-related protein family, even though the GTPase domain is the only conserved sequence. They have various functions in the resistance against intracellular pathogens via GTP binding and hydrolysis [1–5]. Generally, an infection is followed by the production of interferons by leukocytes, monocytes and fibroblasts, leading to transcriptional activation of the interferon-stimulated genes. GBPs belong to the vertebrate specific class of interferon-*γ* induced effector molecules that combat intracellular bacteria, parasites and viruses [6]. The human guanylate binding protein 1 (hGBP1) was found to be involved in the defense against viruses, in particular against the vesicular stomatitis virus and the encephalomyocarditis virus, and bacteria [7, 8]. hGBP1 was also identified as a marker of various cancer types, such as mammary cancer and for cutaneous lupus erythematosus [9].

hGBP1 is a large, multi-domain GTPase with similar, but quite low nucleotide binding affinities for GTP, GDP (guanosine diphosphate) and GMP (guanosine monoposphate) [10]. It can adopt at least two structural states with different binding affinities to partner proteins. The switch between the two functional states is activated by GTP binding, resulting in an ‘active’ state (usually the GTP/GDP-bound form) that binds another hGBP1 molecule leading to dimerization or an effector protein for eliciting the desired effect, and a ‘silent’ state that cannot bind and activate other proteins. The dimerization of hGBP1 occurs through their large GTPase (LG) domains, which stimulates hydrolysis of GTP to GDP and subsequently GMP in two successive cleavage steps [10–14]. The hGBP1 monomer has been shown to be able to also hydrolyze GTP to GDP, but not to GMP [15]. The crystal structure of full-length hGBP1 has been solved in the nucleotide-free, i.e., the apo state (PDB 1DG3) [16] and with the non-hydrolyzable GTP analogue GppNHp bound to it (PDB 1F5N) [17]. In addition, crystal structures of the LG domain monomer with GppNHp (PDB 2BC9) and of the LG domain dimer with GDP/GMP · AlF_3/4_ (PDB 2B8W and 2B92) are available [12].

The hGBP1 structure is divided into three domains as can be seen in Fig 1. The LG domain is the most conserved region from the dynamin family and consists of the first 310 amino acids, structured as an eight-stranded *β*-sheet with six parallel and two antiparallel strands, which is surrounded by six main helices. The GTP-binding site contains four conserved sequence elements G1–G4: the canonical G1 motif or phosphate-binding loop (called G1-P loop henceforth), the G2/switch 1 motif (G2-SW1), the phosphate- and Mg^2+^-binding G3/switch 2 motif (G3-SW2), and the nucleotide-specificity providing G4 motif, which is part of a loop and will be called G4-L2 in the following [18]. This G4-L2 loop is preceded by another loop, which we thus denote as L1. Another key structural element of the G domain is the guanine cap (GC), which forms the protein–protein interface in the hGBP1 dimer [12]. The crystal structures of the nucleotide-free and -bound LG domain suggest that the conformation of the GC goes from an open conformation in apo-hGBP1 to a closed conformation upon GTP binding. The different loops along with their residue ranges and residues key for hydrolysis or dimerization are listed in Table 1 and shown in Fig 1. The second domain is the middle (M) domain (amino acids C311–Q480), which is composed of two two-helix bundles, *α*7/8 and *α*10/11 that are connected by *α*9 and extend over a length of 90 Å, giving hGBP1 an elongated shape. The helical effector (E) domain (amino acids T481–I591) involves a very long helix, *α*12, that stretches over a length of 120 Å from the tip of the M domain back to the LG domain, where it forms multiple electrostatic contacts with helix *α*4’. At the C-terminal end of the E domain, there is a helical turn leading to the short helix *α*13 and the last seven residues, which are unstructured.

**Table 1.** Characterization of the loops of the G domain.

| (Motif-)Loop | Sequence | Key residues<sup>a</sup> | Flexibility |  |  |
| --- | --- | --- | --- | --- | --- |
|  |  |  | clusters | population<sup>b</sup> | RMSD<sup>c</sup> |
| G1-P | 44–52 | K48, K51 | 23 | 89.6% | 7.7 Å |
| G2-SW1 | 58–77 | S73, T75 | 259 | 40.5% | 15.5 Å |
| G3-SW2 | 98–110 | E99 | 175 | 50.6% | 14.5 Å |
| L1 | 149–172 |  | 115 | 42.4% | 9.5 Å |
| G4-L2 | 181–198 | D184 | 194 | 37.1% | 13.4 Å |
| GC | 235–261 | R240, R244, D255 | 675 | 26.4% | 22.0 Å |

<sup>a</sup> Residues which are important for GTP binding, hydrolysis or hGBP1 dimerization.
<sup>b</sup> Percentage of the structures which are cumulatively represented by the first three clusters.
<sup>c</sup> The largest RMSD found between two loop conformations.

**Fig 1.**
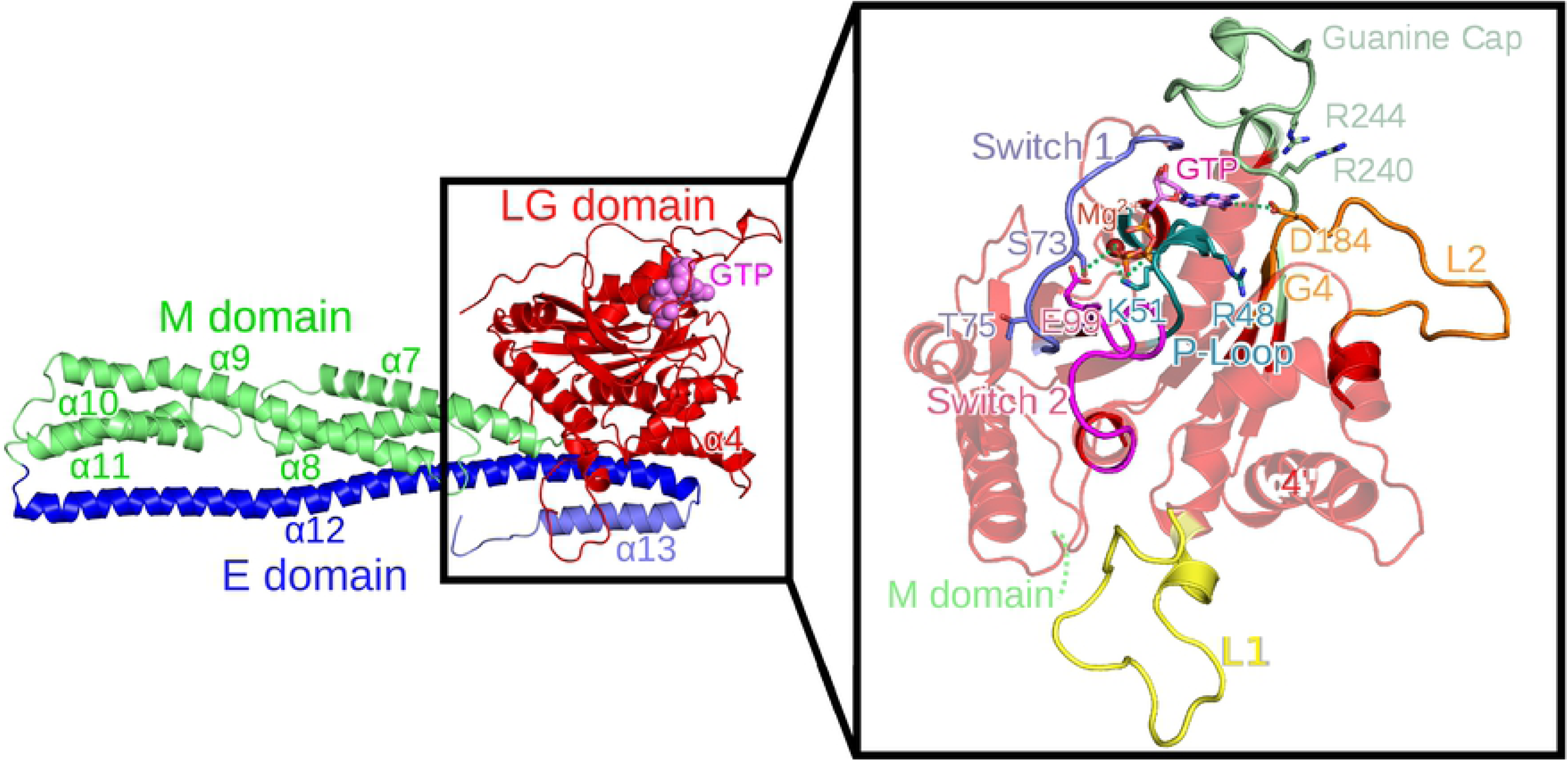
Conformation of the nucleotide-bound hGBP1 crystal structure (PDB 1F5N). The different domains are highlighted in different colors: the LG domain in red with GTP shown in purple, the M domain in green, and the E domain in blue (with different shades used for *α*12 and *α*13 for ease of distinction). Loops missing in the crystal structure were modeled. In the enlargement of the LG domain shown on the right, the four GTP-binding site motifs are displayed: the G1-P loop in turquoise, G2-SW1 in blue, G3-SW2 (G3) in magenta, and G4-L2 in orange. The guanine cap (green), loop L1 (yellow), and residues important for dimerization or GTP binding and hydrolysis (shown as sticks with the same color as the corresponding loop) are also highlighted.

The aim of the current work is to elucidate the intrinsic dynamics of apo-hGBP1 and the hGBP1 dimer. Given the considerable size (67 kDA, 591 residues) and elongated shape of hGBP1, it is to be expected that even without nucleotide binding this protein is flexible. A thorough characterization of the conformational dynamics in the apo state is the prerequisite for understanding the changes in structure and dynamics of hGBP1 following GTP binding, hydrolysis and dimerization. To this end, we applied multi-resolution molecular dynamics (MD) simulations to both the hGBP1 monomer and dimer, involving in total 13 *µ*s of sampling at the atomistic scale and 1.1 ms at the coarse-grained level. To elucidate the dominant motions from the large amount of simulation data, we applied state-of-the-art techniques, such as principal component analysis and Markov state modeling. The enhanced MD simulations of the monomer revealed that the monomeric apo form is highly flexible and exhibits a hinge motion that is similar to the motions observed for other dynamin-like proteins. This motion is also present in the hGBP1 dimer, for which a structural model is provided in this study. Other large-scale motions were observed for the C-terminal helix *α*13, which allows us to explain previously reported experimental data, and for most of the loops of the LG domain loops, which provides a rationale why GTP is required for hGBP1 dimerization to occur. Our study provides fundamental insights into the dynamics of both the hGBP1 monomer and dimer with consequences for its function.

## Results

To reveal the conformational dynamics of apo-hGBP1 on the microsecond time scale, we performed an all-atom Hamiltonian replica exchange MD simulation (H-REMD) with 30 replicas of 400 ns length per replica (see Table 2 in the Methods for an overview of all simulations performed in this study). It should be noted that the usage of the replica exchange algorithm usually leads to a sampling speed-up of one or two orders of magnitude compared to ordinary MD simulations [19, 20]. To explore the dynamics on the sub-millisecond time scale for apo-hGBP1 and also the hGBP1 dimer, we employed coarse-grained MD simulations using the Martini model [21]. We first present the results from the all-atom H-REMD simulation, followed by the results from the Martini simulations.

**Table 2.** List of simulations performed in this work.^*a*^.

| System | Simulation | Size | $T$ | Runs | Length | Cumulated time |
| --- | --- | --- | --- | --- | --- | --- |
| Monomer | Amber99SB*-ILDNP<br>H-REMD | 335,553<br>atoms | 310 K | 1 × 30<br>replicas | 400 ns<br>per replica | 12 μs |
| E domain<br>(T481–I591) | Amber99SB*-ILDNP<br>MD | 573,042<br>atoms | 310 K | 1 | 500 ns | 0.5 μs |
| α11 + E domain<br>(Q456–I591) | Amber99SB*-ILDNP<br>MD | 573,050<br>atoms | 310 K | 1 | 500 ns | 0.5 μs |
| Monomer | Martini<br>MD | 18,438<br>particles | 310 K | 5 | 63 μs, 75 μs, 75 μs<br>75 μs, 200 μs | 488 μs |
| Dimer | Martini<br>MD | 32,836<br>particles | 310 K | 5 | 23 μs, 74 μs, 75 μs<br>90 μs, 150 μs | 412 μs |
| Dimer | Martini<br>MD | 32,836<br>particles | 320 K | 1 | 270 μs | 270 μs |

<sup>a</sup> All simulations reported in this work were performed on the supercomputer JURECA at the Jülich Supercomputing Centre [54].

### Conformational dynamics of the hGBP1 monomer from all-atom simulations

#### Overall flexibility

We evaluated the flexibility of hGBP1 by calculating the root mean square fluctuations (RMSF) of the C_*α*_ atoms. It is found that the LG domain is the most rigid part of hGBP1 while the regions furtherst away from it are the most flexible, which can be best seen in the structure plot in Fig 2 where the rigid amino acids are shown in blue and the mobile parts of the protein are colored in red. The most stable regions are present in the LG domain and involve amino acids that belong to the *α*-helices or the *β*-sheet of that domain. These amino acids were therefore used for aligning the conformations with respect to the initial conformation before calculating the RMSF. The loops of the LG domain can be easily identified as the regions with increased RMSF values (see also Table 1), which will be discussed in detail below.

**Fig 2.**
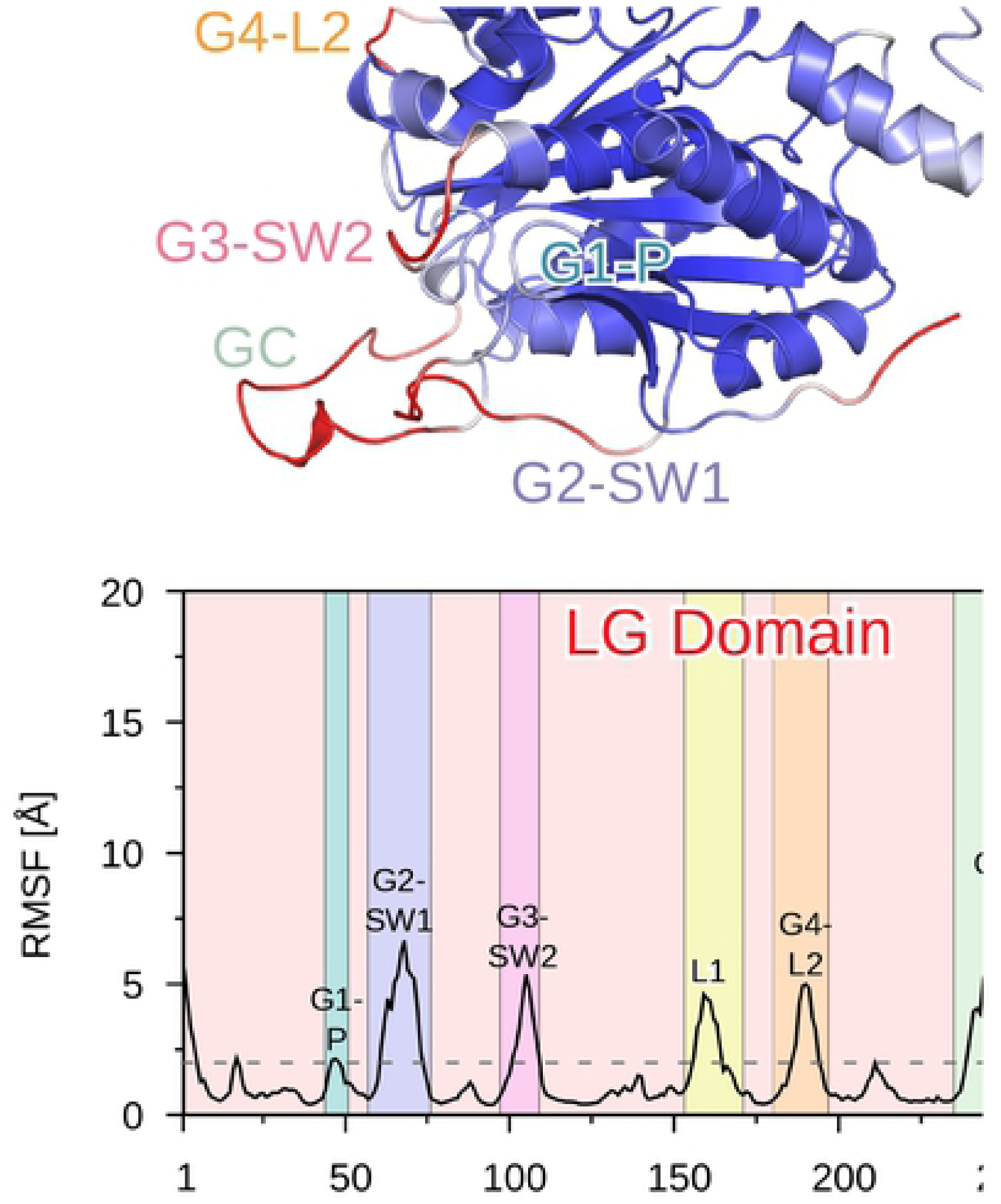
Fluctuations of the hGBP1 residues obtained from the H-REMD simulations. The fluctuations are quantified by the RMSF of the C_*α*_ atoms. The top plot shows the protein colored based on the RMSF values, ranging from blue for low RMSF values (minimum at 0.4 Å) to red for high RMSF values (maximum 15.8 Å). Regions of increased flexibility (loops of the LG domain, regions 1–4 in the M or E domain) are marked in the top and bottom plots. The dashed horizontal line at 2 Å helps distinguishing rigid and mobile regions from each other.

The first highly flexible region of the M domain involves amino acids A320–R370, which form the two-helix bundle *α*7/8. The residues connecting these two helices are the most flexible, resulting in an RMSF peak of ∼7 Å (marked as region 1 in Fig 2). This two-helix bundle is followed by residues L375–T387, which are quite rigid due to their proximity to the LG domain, and the third and longest helix of the M domain, *α*9, which is characterized by monotonically increasing RMSF values up to one of the two highest RMSF peaks of ∼15 Å for residue K429 (region 2 in Fig 2). The next two-helix bundle composed of *α*10 and *α*11 has RMSF values between 7 and 15 Å, with the lowest values found for the residues connecting these two helices (region 3 in Fig 2). The RMSF of ∼7 Å for this turn region is similar to the values for region 2 between *α*7 and *α*8, which can be explained by their close geometric proximity. The second RMSF peak of ∼15 Å corresponds to the transition between domains M and E (region 4 in Fig 2), where the long helix *α*12 from domain E starts. The RMSF values for the residues of this helix decrease until it comes in contact with the LG domain around K544, where the RMSF has dropped below 2 Å. Since *α*13 forms several tight interactions with both *α*12 and the LG domain, this helix was rather rigid in our H-REMD simulation. Only the last seven C-terminal amino acids were flexible, as were the first four N-terminal amino acids.

#### Flexible loops in the LG domain

To characterize the dynamics of the LG loops, we clustered the conformations of each loop using a cutoff of 2.5 Å. The results of this analysis are summarized in Table 1 and shown in Fig 3. Compared to the other loops of the LG domain, the G1-P loop is only slightly flexible. The overall number of clusters is low (23), the first three clusters present most of the loop conformations (89.6%), and the RMSD between the two conformations that are furthest apart is only 7.7 Å, which includes side-chain motions as they were considered during the clustering analysis. Residue R48 is in all clusters solvent-exposed and thus in a position that is not compatible with GTP hydrolysis. This is not too surprising as it is known that only after GTP binding—which was not considered in our simulations—followed by dimerization R48 is positioned toward the *γ*-phosphate of GTP, stimulating the cleavage of this group by stabilizing the transition state of GTP hydrolysis [22]. Interestingly, K51, which is also crucial for GTPase activity of hGBP1, is not flexible and remains in the same position as during GTP hydrolysis.

**Fig 3.**
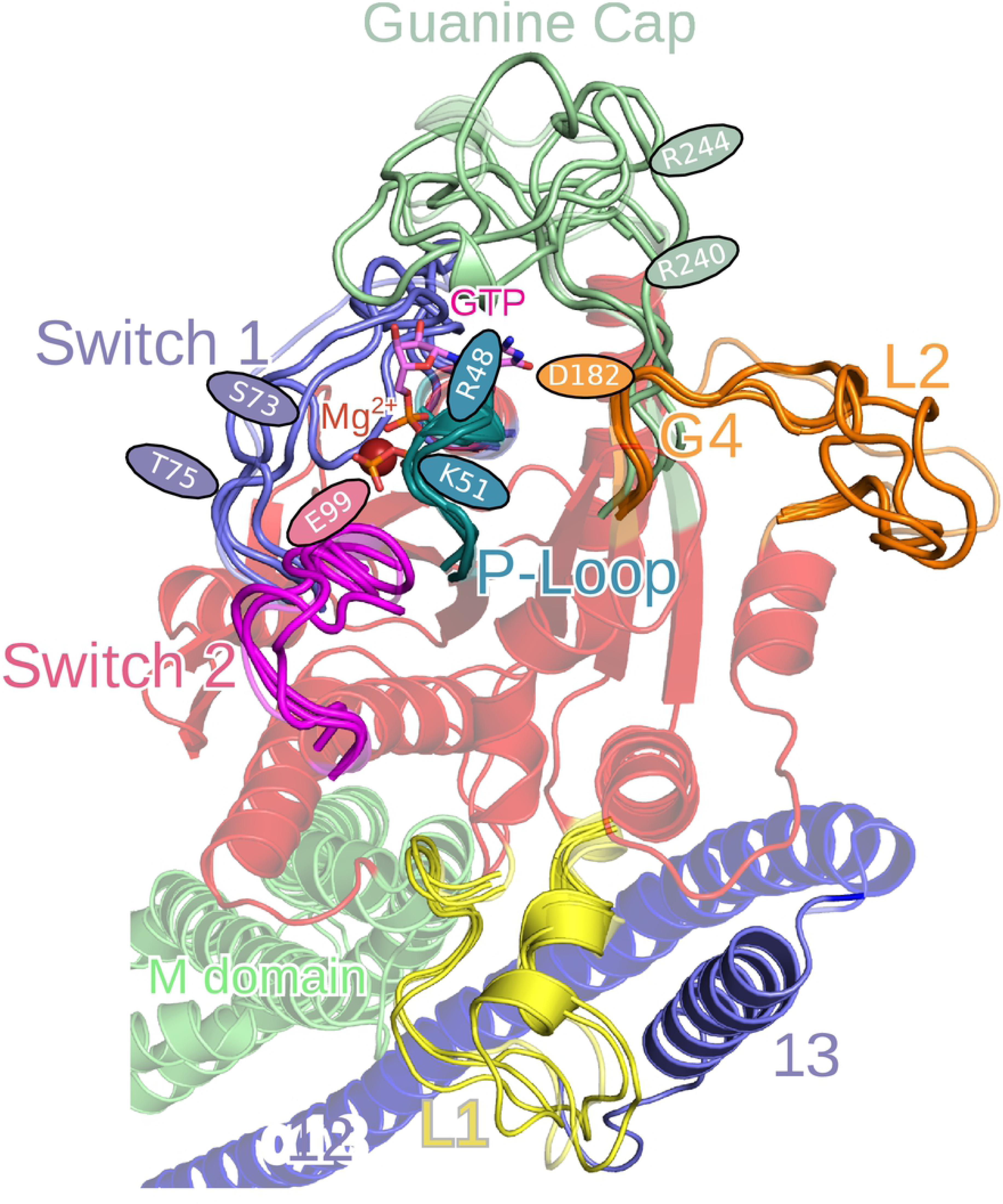
Conformational clusters of the LG domain loops. The LG, M and E domains are colored in transparent red, green and blue, respectively. The colors of the loops are: G1-P in turquoise, G2-SW1 in blue, G3-SW2 in magenta, L1 in yellow, G4-L2 in orange, and the GC in green. The positions of the residues important for GTP binding and hydrolysis or hGBP1 dimerization are indictaed by ellipses in the color of the corresponding loop.

One of the most flexible loops is the G2-SW1 loop, for which 259 clusters were identified. Fig 3 shows that it can switch between closed and open conformations with a preference towards the closed state despite the lack of nucleotide in our simulations. However, the two residues S73 and T75, which are important for the hydrolysis reaction [14, 22], are in positions different from the ones in the LG domain dimer. Without GTP, they point away from the GTP-binding site. Thus, they must undergo a reorientation for adopting positions supporting GTP hydrolysis upon GTP binding and hGBP1 dimerization. In contrast, residue E99, which is part of the G3-SW2 loop and also of relevance for GTP hydrolysis by forming a composite base together with S73, a bridging water molecule, and GTP itself enabling the transfer of a proton from the water nucleophile to the GTP phosphoryl oxygen [14], remains in a position in agreement with GTP hydrolysis. The stable orientation of the E99 side chain allows its interaction with the Mg^2+^ ion (which was not present in our simulations), while residues 101–110 of G3-SW2 are flexible. Loop L1, which is not part of the GTP binding site or dimerization interface, is particularly flexible between residues 156 and 165, while the other residues remain in their position. This is understandable as some of these residues belong to an *α*-helix (see Fig 3), but we nonetheless included them in our analysis because they interact with helix *α*13 from domain E. We found that the interplay between the L1–*α*13 interactions and the flexibility of L1 is essential for enabling large-scale motions of *α*13, which is discussed below. The G4 motif, which contains D184 relevant for GTP hydrolysis, is stable as it is part of a *β*-strand, while the following loop L2 is very flexible with motions of up to 13.7 Å and 194 clusters in total. The firm position that is found for D184 enables efficient binding of the guanine base of GTP to this residue.

The highest flexibility—with 675 clusters and the first three clusters representing only 26.4% of the conformations—was found for the GC loop relevant for hGBP1 dimerization. Moreover, large confomational changes are possible as the maximum RMSD of 22.0 Å revealed and can be seen in Fig 3. Both open and closed conformations are adopted by the GC with many different conformations between these extremes, which agrees with the findings for other enzymes that loop motion is often not a simple open and shut case [23]. The presence of the closed GC state without GTP being bound and hGBP1 being dimerized further suggests that GTP only helps stabilizing the closed GC state and does not induce a conformational transition from the open to the closed state, as one might assume from the crystal structures [12, 16].

#### Motions of the M domain and and helix *α***12**

The main structural changes of the M domain and helix *α*12 of the E domain were identified based on a principal component analysis (PCA) of the H-REMD data (see Methods for details). Helix *α*13 was excluded from this analysis as the RMSF values had revealed that it did not substantially move. We found that the first two principal components (PCs) describe best the main conformational motions of the M domain and *α*12 (Fig S1 in the Supplementary Information) and therefore calculated the 2D free energy surface (FES) of hGBP1 along these two PCs (Fig 4). The main conformational change along the PC1 is a kinking motion, were the M domain and helix *α*12 bend towards the LG domain for negative values and away from it for positive values. The amplitude of this motion is large as the distance between the minimum and maximum values for this motion is above 110 Å, which becomes visible from the conformations corresponding to PC1_min_ and PC1_max_ in Fig 4. The motion along PC2 can be described as a screwing motion where the M domain and *α*12 rotate with respect to the LG domain, which can be seen by comparing the conformations representing PC2_min_ and PC2_max_. In general, the FES is characterized by one main area which contains the lowest free energy minimum. Conformations corresponding to the lowest free energy values (shown on the right of Fig 4) have structures that are very similar to the crystal structure [12, 16]. In addition, there is a shallow free energy minimum for negative values of PC1 that corresponds to conformations where the M domain and helix *α*12 are bent towards the LG domain. Interestingly, in the structure representative for this minimum the long helix *α*12 from the E domain is broken into two shorter helices. A similar helix fragmentation, which preferentially occurred around residue Q541, is also seen in the structures representing PC1_min_ and PC2_min_.

**Fig 4.**
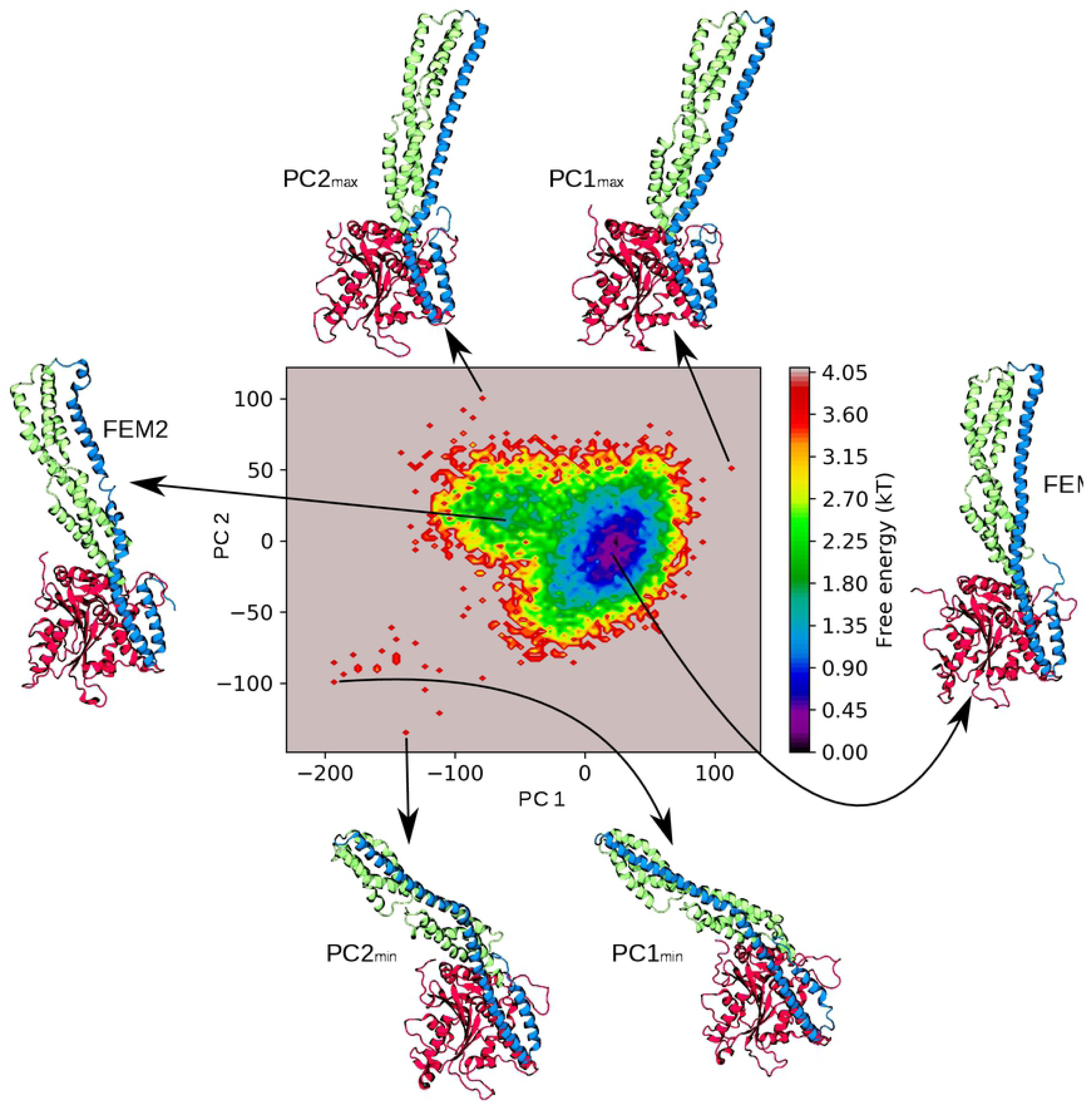
Free energy surface (FES) along the dominant principal components PC1 and PC2. Next to the FES, hGBP1 conformations representing the lowest free energy minimum (FEM1, right) and the shallow free energy minimum (FEM2, left) are shown. Examples of structures for the extreme values along PC1 and PC2 are also provided below (PC1_min_, PC2_min_) and above (PC1_max_, PC2_max_) the FES. The conformations are shown as cartoon and colored based on the three domains: LG (red), M (green) and E (blue).

Despite these large-scale motions of the E domain, it did not detach from the LG or the M domain due to the presence of multiple stable salt bridges formed by the E domain with both other domains (Fig 5A and B). The most stable salt bridge was present between K228 from helix *α*4’ of the LG domain with either E568 or E575 of *α*13, which existed during 43.0% of the simulation time. Other residues with a high likelihood of salt-bridge formation are K582 and K587 of *α*13, which built salt bridges with various residues of the loop L1 of the LG domain, with a probability of 43.0% and 7.1%, respectively. The decrease of the electrostatic interactions as done in our H-REMD simulations did not completely abolish these salt bridges, confirming their stabilities. In the replica with the largest energy bias, residues K228, K582 and K587 formed the same salt bridges as in the target replica with probabilities of 26.2%, 18.0% and 1.3%, respectively, which was still enough to inhibit the detachment of the E domain from the LG domain.

**Fig 5.**
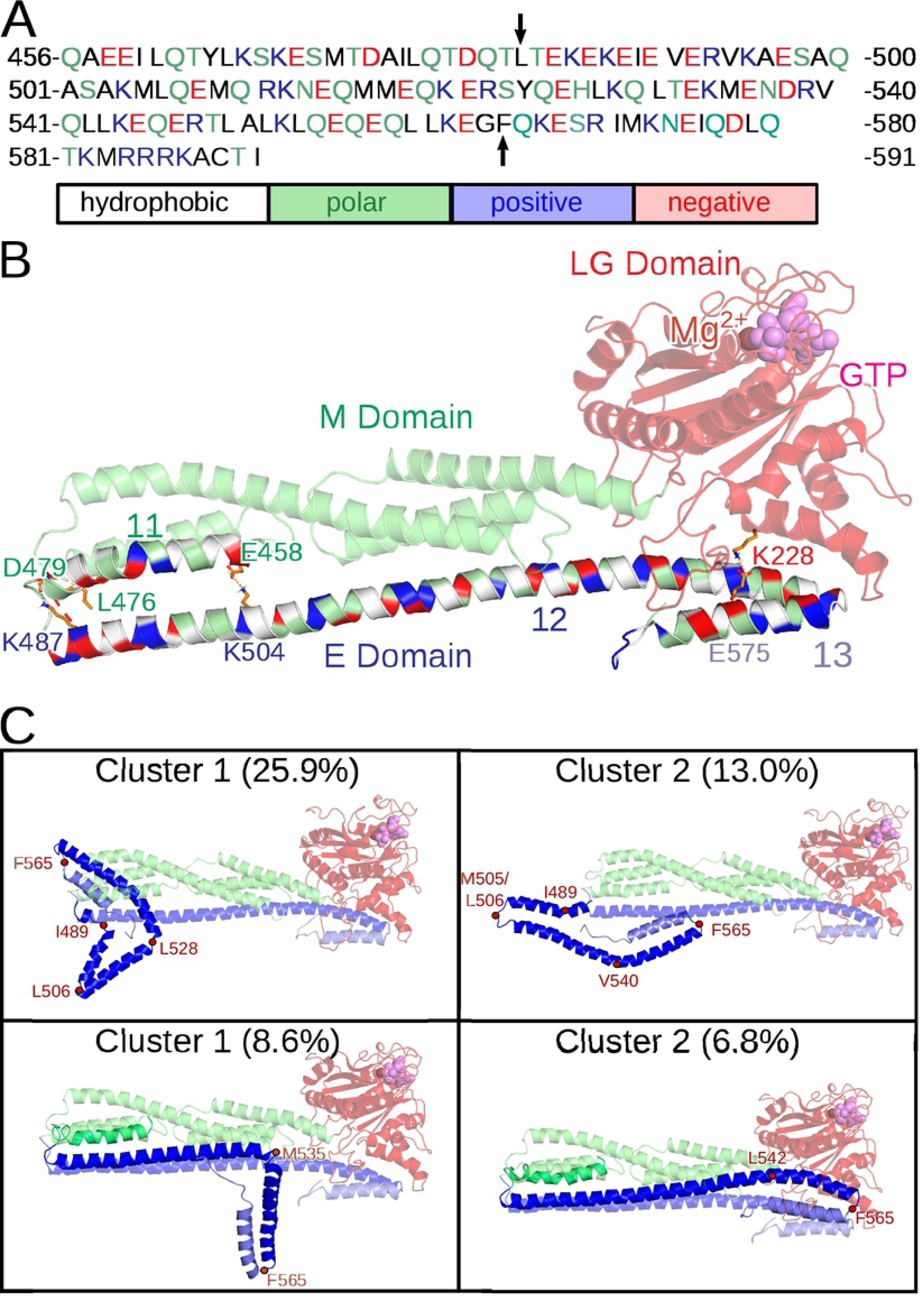
Properties and dynamics of the isolated E domain of hGBP1. (A) The sequence of the E domain and preceding helix *α*11 of the M domain is shown and colored according to the physicochemical properties of the amino acids. The start and end of helix *α*12 at residues L482 and G564 are indicated by vertical arrows. (B) The structure of the E domain and its interactions with the LG domain (red transparent) and the helix *α*11 of the M domain (green transparent) are highlighted for the crystal structure of hGBP1 (PDB 1DG3). The residues of *α*11 and the E domain are colored according to the physicochemical properties of the amino acids: white, hydrophobic; green, polar; blue, positively charged; red, negatively charged. Salt bridges (K228–E575, E458–K504, D479–K487) are indicated with orange dashes and hydrogen bonds (L476/D479–K487) with black dashes. The interacting residues are shown as sticks and labeled. This inludes K582 and K587, which form salt bridges with various residues of loop L1 (in yellow) of the LG domain. (C) The two most populated cluster conformations with their occurrence obtained from the MD simulations of the isolated E domain (top) and of *α*11 plus E domain (bottom). The cluster structures were aligned to the crystal structure of full-length hGBP1 (shown as transparent cartoon) using residues 482–484 for the alignment of the isolated E domain and *α*11 for alignment of *α*11 plus E domain. The red points indicate the turns that formed at the different positions of the E domain along with the turn at G564 separating *α*12 and *α*13 from each other.

#### Dynamics of the isolated E domain

Motivated by the dynamics of the E domain observed in our H-REMD simulation and by the hypothesis, derived from size-exclusion chromatography, that hGBP1 can adopt an even more elongated shape than it already has by folding out the E domain (see Fig 7 in [24]), we performed 500 ns MD simulations of the isolated E domain and also of the E domain plus helix *α*11 of the M domain (see Fig 5A for the sequence). These simulations allow us to judge the stability of the E domain when it is not stabilized by intraprotein interactions with the LG and M domains. While the lack of these tertiary contacts would be the prerequisite for folding out of the E domain, our simulations revealed that the E domain is not stable without them. In the simulation of the isolated E domain helix *α*12 formed kinks and turns at different positions, especially at the N-terminal end, which in hGBP1 is attached to the M domain. In Fig 5C the two dominant structures as obtained from a clustering analysis of the 500 ns MD simulation are shown (structures for the next three clusters are shown in Fig S2A). We overlaid the E domain conformations onto the crystal structure of hGBP1, which demonstrates that the E domain is unlikely to fold out as an extended helix if it should detach from the LG domain, as suggested from experiments [24]. The two most stable kinks or turns occured at residues I489 and M505/L506, while the region between Q525 and L542 transformed to turns at various positions, but always reversibly as can be seen in the time-resolved secondary structure plot in Fig S2B. The latter position is the same where *α*12 reversibly unfolded and kinked during the H-REMD simulation of full-length hGBP1 (Fig 4).

**Fig 6.**
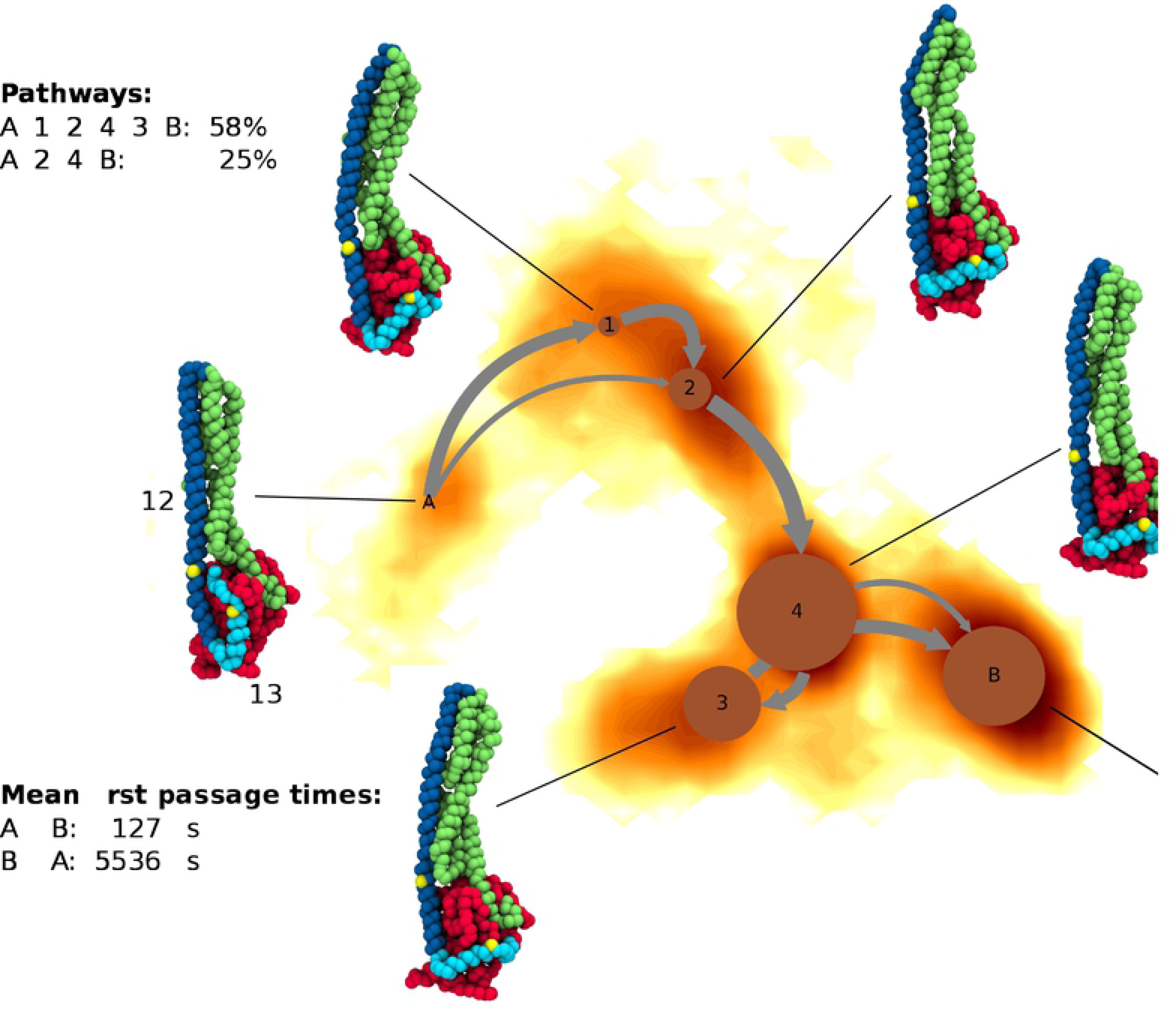
A network diagram of the Markov state model obtained for the motion of helix *α*13 as sampled in the Martini simulations. The MSM is overlaid onto the FES along the first two TICs. The circles represent the stable states, where the area of the circles correlates i with the population of the corresponding state. The arrows indicate transitions between the states, with the line thickness correlating with the transition fluxes. The FES is colored as gradient from yellow (high free energies) to dark red (low free energies). A representative conformation is shown for each state and colored based on the three domains, while the yellow dots indicate residues Q541 and T581, which are the start and end of the sequence for which the MSM was calculated.

**Fig 7.**
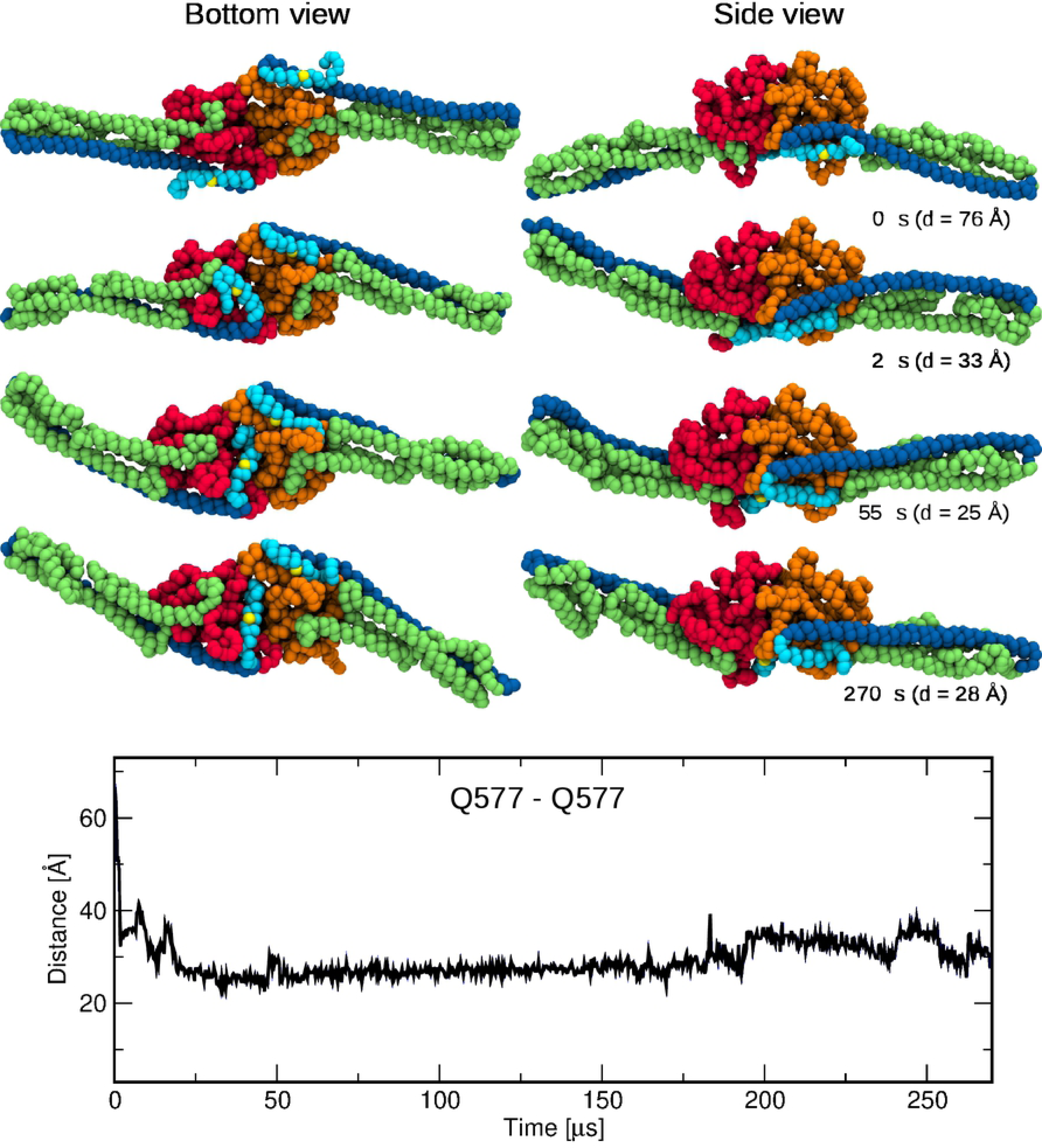
Motion of helix *α*13 in the hGBP1 dimer. This motion is monitored by the time evolution of the distance between the two residues Q577 from *α*13 of the two proteins composing the dimer. Representative snapshots associated with different times are also shown. The two LG domains are shown in red and orange, respectively, for ease of distinction, while the M domains are shown in green and the E domains in blue. The two Q577 residues are indicated by yellow spheres.

We aimed at understanding the reasons for turn formation that occurred in the isolated *α*12 helix. This helix is 81 residues long and contains 21 *i, i* + 3 or *i, i* + 4 combinations of oppositely charged side chains (Fig 5A), which may form salt bridges. While such salt bridges are known to stabilize helices [25, 26], we find that in some cases they can also stabilize turn structures. For instance, the interaction E526–K529–E533, which is present in *α*12, also stabilizes the turn formed at L528 in the second half of the MD trajectoy of the isolated *α*12 helix. Other salt bridges were newly formed upon turn formation, such as the one between E488 and R493 following the helix → turn transition at I489. In the helix conformation, E488 cannot form a salt bridge with residues 491 or 492 as these are valine and glutamate. Also the turn at M505/L506 enabled a novel salt bridge to be formed, namely between E508 and R511, which in the helix is not possible as the side chains of these two residues point in opposite directions. Also between E508 and K504 only a weak interaction is possible in *α*12 due to steric constraints of the helix (the minimum distance between both side chains is above 6 Å), while upon turn formation E508 becomes able to interact with R511 instead (minimum distance below 2 Å). Interestingly, the predicted positions of turn formation coincide with the positions with a reduced *α*-helical coiled-coil probability of the E domain, which was determined by Syguda et al. [27] using the the COILS program [28]. They performed this analysis to test whether coiled-coil formation may occur between *α*12 helices. An alternative scenario would be that the isolated E domain forms an intraprotein coiled-coil structure with the first helix extending from L482 to A503 and a second helix from L506 to the end of *α*12. A third helix with a turn around residue 540 might also be possible, which would be in agreement with both our MD simulations and the COILS predictions. It should be further noted that the helical propensity of the isolated E domain is only slightly reduced upon turn formation. It is 79% averaged over the 500 ns MD simulation, which is close to the value of 82% extracted from the circular dichroism (CD) spectrum of the E domain [27], especially if one considers the errors involved in fitting CD spectra. In the crystal structure (PDB 1DG3) the helical probability is 86% for *α*12/*α*13, while it was 84% in the H-REMD simulation of hGBP1. Thus, the presence of the LG and M domains have stabilizing effects on *α*12, but the overall helicity is not lost.

A major stabilizing effect originates from *α*11 of the M domain, as its inclusion in the simulation of the E domain prevented the formation of turns at residues I489 and M505/L506. This can be explained by attractive interactions between *α*11 and *α*12, inhibiting structural changes in this part of *α*12. Especially residues close to the turns that formed in the isolated E domain are involved in interactions with *α*11: two stable hydrogen bonds are present between L476–K487 and D479–K487, supported by a salt bridge between E458 and K504 (Fig 5B). Only at different positions between Q525 and L542 reversible turn formation was observed as the cluster structures and evolution of the secondary structures confirm (Figs 5C and S2). For instance, in the most populated cluster structure a turn at M535 is present, which is facilitated by a novel electrostatic interaction between D538 and R539, while preserving the salt bridge E536–R539 present in *α*12. In both simulations, i.e., in the simulation of the isolated E domain and in that of *α*11 plus E domain, *α*12 tightly interacted with *α*13 and the last seven C-terminal residues via various salt bridges, which are possible as 11 of the 28 amino acids composing *α*13 and the C-terminal residues are charged (including the stretch R584-R585-R586-K587) and paired with a similar abundance of positively and negatively charged residues in the oppositely located *α*12 helix.

In summary, the simulations of full-length hGBP1, the isolated E domain, and *α*11 plus E domain revealed that a folding out of the E domain seems to be an unlikely event. First, it would require the simultaneous breaking of several hydrogen bonds and salt bridges that the E domain forms with both the LG domain and the M domain. Second, even if such detachment of the E domain should happen, the missing tertiary contacts would provoke turn formation at specific residues of *α*12 on the nanosecond time scale, as demonstrated by our MD simulations. It should also be mentioned that this helix is by a factor of at least four longer than the optimal length of an isolated helix (which is between 9 and 17 amino acids [29]) and thus needs tertiary contacts as present in hGBP1 for it to be stable. Instead of folding out of *α*12 as an extended helix, a more likely conformational change might be the formation of a coiled-coil motif following fragmentation of *α*12 into two or three shorter helices. In all cases studied here, *α*12 has a high tendency of reversible kinking around residues Q525–L542. Even in full-length hGBP1 this kinking is possible, as also the M domain is very flexible in that area, which is where the two-helix bundles *α*7/8 and *α*10/11 meet (marked as regions 1 and 3 in Fig 2). We thus conclude that the M and E domains feature a hinge in that area.

### Long time-scale dynamics of the hGBP1 monomer and dimer from coarse-grained simulations

#### Monomer dynamics

To explore the conformational flexibility of the apo-hGBP1 monomer on the micro- to millisecond time scale, we performed five continuous MD simulations of lengths between 63 *µ*s to 200 *µ*s using the coarse-grained Martini force field (see Table 2). As in the all-atom simulations we observed the kinking motion of the M and E domains in all five coarse-grained simulations. Interestingly, this motion was possible despite the fact that the Martini model preserves the initial secondary structure by applying an elastic network model. Therefore, *α*12 remained fully helical in these simulations. It further demonstrates that the application of Martini without elastic networks to preserve the tertiary protein structure allows the sampling of conformational changes that also occur in atomistic simulations.

Another substantial motion observed in the Martini simulations was a change in orientation of helix *α*13 by ∼90° with respect to helix *α*12. To describe this motion we applied a Markov state model (MSM) analysis to amino acids Q541–T581, which contain the last third of *α*12 and the helix *α*13 (see Methods and Figs. S3 and S4 for details of this analysis). We identified six metastable states, for which representative conformations are displayed in Fig 6 together with the MSM overlaid onto the FES along the first two time-lagged independent components (TICs). Since all simulations started from the crystal structure, we labeled the Markov state that had ∼40% of the conformations with helix *α*13 close to helix *α*12 and which are thus similar to the crystal structure with A. The state corresponding to the largest displacement of *α*13 is denoted as state B, while the other four Markov states (labeled 1–4) are intermediate states between A and B. We calculated the reactive fluxes between the states, which are shown as gray arrows in Fig 6, and determined the mean first passage time (MFPT) needed for the system to go from state A to state B, obtaining a value of 127 *µ*s. The MFPT for the reverse transition from B to A is 5,536 *µ*s, which is consistent with the observation that the complete reverse motion of helix *α*13 to the conformation associated with the crystal structure has not occurred in our simulations. It should be noted that the MFPTs reported here are obtained from coarse-grained simulations and might not be equivalent to actual MFPTs. Generally, events observed in simulations with the Martini force field are between 3 to 5 time faster than similar events observed in atomistic simulations [21]. The main transition pathway from A to B goes via states 1, 2, 4, and 3 with a probability of 58%. The next most significant transition pathway with a probability of 25% involves only two intermediate states, 2 and 4.

The mobility of helix *α*13 on the surface of the LG domain requires its motion beyond the loop L1 from the LG domain. Our H-REMD simulations had revealed that especially K582 and K587 have a high tendency to form salt bridges with different residues from that loop (D159, E160, E162, E164, D167, D170, see Fig 5B). In order to identify the molecular interactions enabling the motion of *α*13 beyond this loop, we investigated the interactions between the C-terminal region F565–T590, which includes *α*13 (F565–M583), and L1 by calculating an average distance map during the simulation interval when this motion occurred. The distance map (Fig S5A) revealed that the strongest contacts are between the sequence ^580^QTKMRRRKA^588^ of the C-terminus and the sequence ^164^EVEDSAD^170^ from L1. This indicates that the motion of *α*13 is facilitated by electrostatic interactions in interplay with the high mobility of the loop L1 (see Fig 3), allowing the C-terminal region to move beyond L1 and then further on the surface of the LG domain. We also investigated the cause for the high stability of *α*13 after it has moved beyond L1, yielding Markov states 3, 4 and B, and a very large MFPT for the reverse transition from B to A. To this end, we analyzed the interactions between the C-terminal region F565–T590 and the LG domain for conformations belonging to Markov state 4, which has the largest population. The average distance map between the C-terminal region and the interacting LG domain region is shown in Fig S5B, with two notable interactions involving helices from the LG domain being highlighted. In the first case, sequence Y144–K155 belonging to helix *α*3 is in contact with the second part of *α*13 (I576–M583) and the following, flexible residues R584–A588. The strongest contacts are formed between polar residues: E146 and T149–R151 from the LG domain and Q580–T581 from *α*13. The second interaction hot spot involves sequence N220–F230, i.e., half of helix *α*4’ that extends from S213 to F230, which is in contact with residues S569–Q580 from the first part of *α*13. Here, the strongest contacts are mainly of hydrophobic nature, involving L224 and C225 from the LG domain and M572 and I576 from *α*13. In addition, helix *α*13 has residue L579 exposed to the LG domain, constituting another hydrophobic interaction. Thus, we conclude that the initial motion of *α*13 is driven by the coaction of the electrostatic interactions between *α*13 and L1 and the flexibility of L1, and once the helix has moved beyond this loop, it is stabilized by hydrophobic and polar interactions with helices *α*3 and *α*4’ from the LG domain.

#### Dimer model from coarse-grained simulations

The hGBP1 dimer is formed between GTP/GDP-bound monomers and is thought to be the biologically active state of the protein [13]. Starting from the crystal structure for the LG domain dimer of hGBP1 (PDB 2B92) [12] and the crystal structure of the hGBP1 monomer (PDB 1DG3) [16] we built a complete hGBP1 dimer model as described in the Methods section. We converted the atomistic conformation to a coarse-grained Martini model and performed five simulations of different time lengths (between 23 *µ*s and 150 *µ*s, see Table 2). In addition to restraining the *β*-sheet and the helices of the LG domain to their original positions (see Methods), the residues of the G2-SW1 and GC loops were also restrained. They thus remained in a position of the transition state during GTP hydrolysis as GDP ·AlF_3_, which is present in the PDB structure 2B92, is a transition-state mimic. Our Martini simulations of the hGBP1 dimer can thus be considered to represent the nucleotide-bound state even though the nucleotide was not present during the simulations.

As for the monomer, we observed the kinking motion of the M and E domains in all five dimer simulations. However, the change in orientation of helix *α*13 as seen in the coarse-grained simulations of the monomer did not occur. It remained in close proximity to helix *α*12. Only when the temperature was increased to 320 K, as done for one simulation of 270 *µ*s length (see Table 2), we observed this conformational change, but only in one of the proteins composing the hGBP1 dimer. This indicates that the energy barrier for the this motion must be higher than in the monomer. To find the cause for this, we analyzed whether the interactions between *α*13 and loop L1 or whether the mobility of that loop would be different in the dimer than in the monomer. Though both cases did not apply. Instead we found that a new salt bridge between *α*4’ and *α*13 involving residues E217 and K567 is formed in the dimer, which became possible by the somewhat different position that *α*4’ adopts in the dimer than in the monomer: it is by ∼6 Å further away from the core of the LG domain, bringing this helix closer to *α*13 (Fig S6A). In each of the dimer simulations at 310 K this salt bridge stayed intact, thereby preventing the 90° motion of *α*13. At 320 K, on the other hand, in the protein of the dimer withe the flexible *α*13 helix this salt bridge broke (Fig S7).

To describe the motion of helix *α*13 that was observed in the simulation at 320 K, we monitored the distance between the residues Q577 of *α*13 from the two proteins (Fig 7). As can be seen from the time evolution of the distance and the snapshots associated with different times, *α*13 in one of the proteins (the one with the LG domain shown in red) has moved beyond the corresponding loop L1 within 2 *µ*s, and after 55 *µ*s it has adopted a position similar to the one in state B identified in the monomer simulation (Fig 6), where it stayed until the end of the 270 *µ*s simulation. Helix *α*13 from the other protein of the dimer remained in close contact to *α*12 throughout the whole simulation. Nevertheless, as shown in Fig 7, the distance between the two Q577 residues decreased from its initial value of 76 Å to values of ∼30 Å, which is very similar to the values between 22 and 35 Å reported from double electron–electron resonance (DEER) and Förster resonance energy transfer (FRET) studies of the hGBP1 dimer [30], despite the fact that the experimental distances refer to distances between DEER spin labels and FRET dye labels, respectively. The more important aspect is that both experimental techniques predict a change in the Q577–Q577 distance by 50% (FRET) or even more (DEER) upon dimer formation compared to the distance that one obtains from the dimer model that is built based on the crystal structure of the hGBP1 monomer.

## Discussion

We studied the conformational dynamics of the nucleotide-free hGBP1 monomer and the hGBP1 dimer using multi-resolution and enhanced MD simulations on the micro- to millisecond time scale. As expected from its highly conserved sequence in the dynamin family, the LG domain is overall very stable in our atomistic H-REMD simulation. The residues involved in the *β*-sheet and the adjacent *α*-helices have all fluctuations of less than 2 Å, while apart from one case the LG domain loops were found to be flexible (Fig 3 and Table 1). The less flexible loop is the phosphate-binding loop G1-P and involves the first of the four motifs G1–G4 that are important for the hydrolysis reaction. These motifs include six residues (R48, K51, S73, T75, E99 and D184) that are directly or indirectly involved in GTP binding and hydrolysis. Even without nucleotide being present, K51, E99 and D184 adopt the same and stable orientations as in the crystal structures of hGBP1 and the LG domain with nucleotide being bound, while the other three residues are flexible without the stabilizing interactions with GTP. This suggests that the active site of the GTPase domain is quite flexible compared to those of other enzymes [31] and only partially preorganized prior substrate binding [32, 33], which might explain the rather low binding affinity of hGBP1 for GTP (*K*_*m*_ = 470 *µ*M) [10]. The LG domain loop with the highest flexibility is the guanine cap (GC), which forms the protein–protein interface in the hGBP1 dimer. Even without nucleotide, the GC can adopt both open and closed conformations and rapidly switch between them. This finding indicates that GTP binding would shift the equilibrium toward the closed GC state, which in turn would facilitate hGBP1 dimer formation via the GC–GC interface requiring a certain stability of the protein recognition motif. Dimer formation, in turn, would further stabilize the closed GC conformation and also the orientations of R48, S73 and T75 in positions supporting GTP hydrolysis, which would explain why the hGBP1 dimer is better able to hydrolyze GTP than the hGBP1 monomer is. Our future atomistic simulations of the GTP-bound hGBP1 monomer and dimer will address whether this hypothesis holds.

One of the main results from our atomistic and coarse-grained simulations is the highly flexible nature of the M domain and the long helix *α*12 from domain E. The region of highest structural flexibility was found at the middle of the M and E domains by PCA (Fig 4), giving rise to large-scale kinking and screwing motions performed by both domains, which is evident from large RMSF values of about 15 Å at the tip of both domains (Fig 2). During these motions, the E domain remains thethered at its ends to both the M and the LG domain by several salt bridges, while the middle of the long helix *α*12 can reversibly unfold and fold, allowing its kinking and screwing. This finding is further supported by the MD simulations of the isolated E domain (with and without *α*11 from the M domain), which also revealed reversible turn formation accompanied with local unfolding between residues Q525 and L542, while the contacts to the M domain were found to be vital for overall stability of the long helix *α*12. Without these tertiary interactions, *α*12 unfolds on the nanosecond time scale, making it unlikely that the ∼120 Å long E domain folds out as intact helix [24], which in addition would require the combined breaking of several hydrogen bonds and salt bridges that the E domain forms with the LG and M domains.

An alternative scenario is a motion similar to that of the bacterial dynamin-like protein (BDLP) from *Nostic punctiforme* (see Fig 8A for a schematic of this motion), which was shown to exist in a closed and extended conformation (PDB codes 2J69 and 2W6D, respectively) [34, 35]. In Fig 8B the closed and extended conformations of BDLP can be seen, along with the most stable hGBP1 structure and the one with the maximal motion of the M and E domains sampled in our H-REMD simulations (corresponding to FEM1 and PC1_min_ in Fig 4) in Fig 8C. The motions of BDLP are facilitated by two hinge regions. Hinge 1 separates the long tail of BDLP into a neck and trunk region, while hinge 2 is at the interface between the neck and G domain. The transition between these two BDLP conformations, which occurs upon nucleotide and lipid binding, involves a 135° kinking between the neck and trunk around hinge 1 and a 75° rotation of the G domain around hinge 2 [34]. In hGBP1 we identified hinge 1 in the region encompassing residues Q525–L542 of the E domain and the meeting point between the helix-bundles *α*7/8 and *α*10/11 of the M domain. It enables a kinking motion of these two domains and involves a (reversible) unfolding of *α*12, dividing it into a short helix close to the LG domain, which would correspond to the neck of BDLP, and a longer helix corresponding to the trunk of BDLP. The presence of a hinge 2 in hGBP1 remains to be shown. Another similarity between BDLP and hGBP1 is that the tip of the trunk of BDLP is the region of highest flexibility.

**Fig 8.**
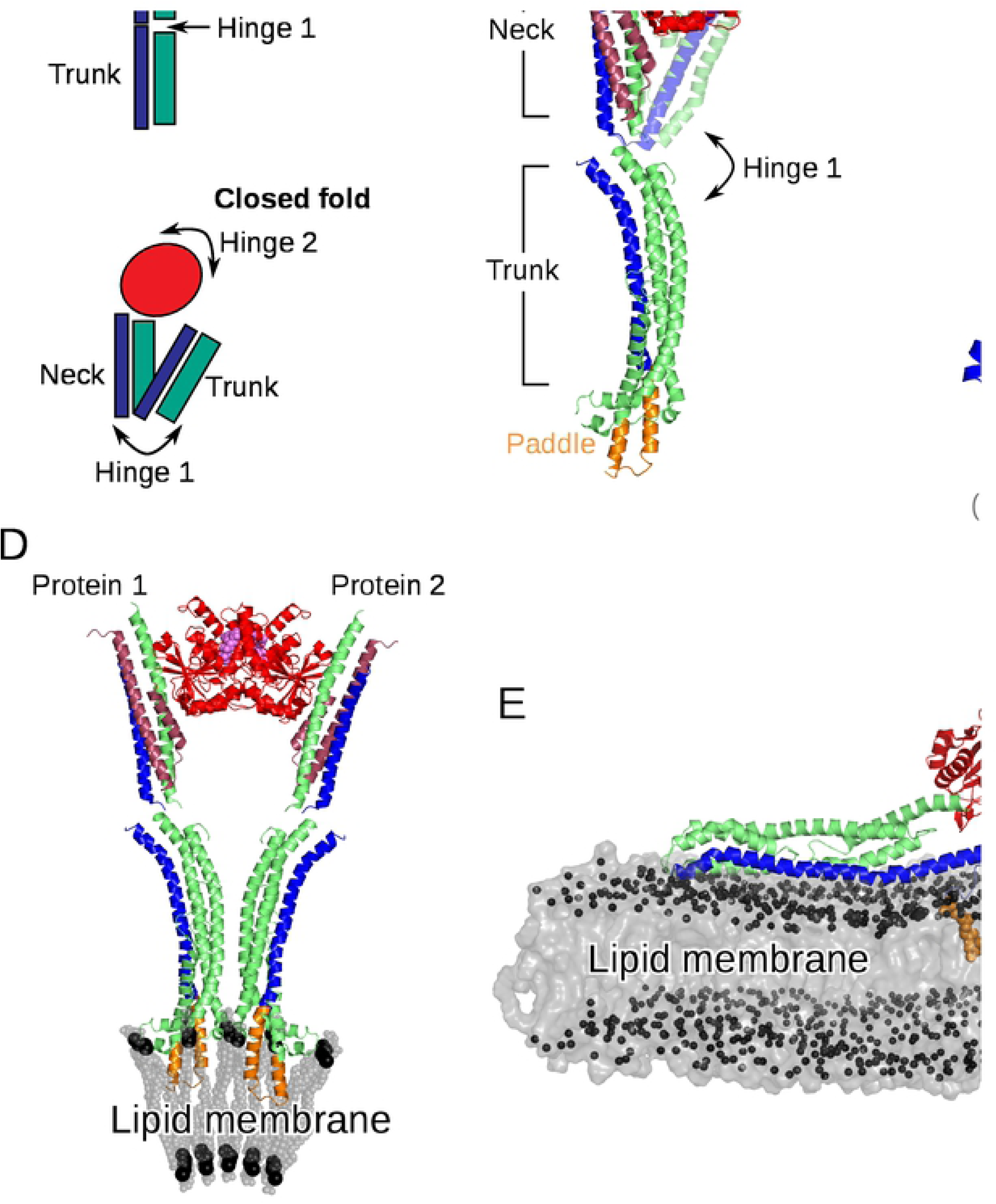
Conformational transitions of hGBP1 and a bacterial dynamin-like protein (BDLP) of *Nostoc punctiforme*. (A) Schematic providing nomenclature for the open and closed BDLP conformations with large-scale transitions between them mediated by hinges 1 and 2 at the trunk/neck and neck/G-domain interfaces. As for hGBP1, the GTPase domain is colored in red with the position of the nucleotide in the open conformation indicated by a magenta sphere, while the 4-helix bundel composing the neck and trunk is colored in green and blue. (B) The closed conformation of BDLP was solved by X-ray crystallography (PDB 2J69, shown in transparent cartoon), while the open conformation was determined by cryo-electron microscopy (PDB 2W6D, shown in opaque cartoon). The two helices preceding the G domain are ruby-colored. (C) From our MD simulations of hGBP1 we observed a kinking of the M and E domains resembling the initial stages of the closed ↔ open transition of BDLP around hinge1, as the overlay of the structures representing this motion (i.e., FEM1 and PC1_min_ in Fig 4) shows. For comparison the crystal structure of hGBP1 (PDB 1DG3) is also shown (in gray). (D) Membrane binding of BDLP occurs following nucleotide binding via the paddle domain (shown in orange) in the open conformation and aggregated BDLP state. Aggregation involves association of the G domains. (E) Membrane binding of hGBP1 also requires nucleotide binding and is facilitated via farsenylation (shown in orange) at the C-terminus acting as lipid anchor.

It should be noted that there are also certain differences between the two proteins. First, the closed state is the preferred conformation of BDLP in its apo form [34], while for hGBP1 the crystal structures show that the extended conformation exists for both the nucleotide-free and -bound form. In fact, at the moment there is no experimental evidence for a closed form of hGBP1. On the other hand, BDLP is not the only dynamin-like protein for which a hinge 1 motion has been revealed. A recent FRET study combined with X-ray crystallography of the human myxovirus resistance protein 1 (MxA) revealed that also this dynamin-like protein can adopt a closed conformation in addition to the open crystal structure [36]. In contrast to BDLP, the open MxA conformation is preferred in its nucleotide-free state, while adding of GTP shifts the equilibrium towards the closed state. The findings for MxA and our results in addition to the structures for BDLP raise the likelihood for a general existence of a closed state for dynamin-like proteins, which should be addressed in the future.

Another difference between hGBP1 and BDLP is the presence of a paddle domain in BDLP at the trunk tip via which membrane binding is facilitaed, while hGBP1 binds to the membrane following farnesylation at the C-terminus (see Fig 8D and E, respectively). This suggests different membrane binding mechanisms for both proteins. Moreover, also the multimerization seems to be faciliated via different domain interactions in BDLP and hGBP1. The dimer structure of the LG domain obtained by X-ray crystallography [12] implies an elongated geometry with the M and E domains pointing away from the dimer interface (Fig 7), leading to a slightly curved dimer which might induce membrane bending following membrane binding. Such membrane destabilization might be even more likely as a result of the kinking motions of the M and E domains observed in our study. In contrast, multimerization of membrane-bound BDLP is thought to occur via interactions between neighboring neck and trunk helices and a G domain dimer interphase different to that seen for the LG domain dimer of hGBP1 (Fig 8D). Our future simulations will address hGBP1 motions following GTP binding and hydrolysis, its multimerization and lipid binding, which will further highlight possible differences and similarities to other dynamin-like proteins, such as BDLP and MxA.

In the coarse-grained simulations of both the hGBP1 monomer and dimer, an important conformational change observed was the change in orientation of the short helix *α*13 of the E domain. Markov state modeling indicated an MFPT for the complete *α*13 motion of ∼127 *µ*s, while the reverse motion is more than an order of magnitude less likely. The slow time scale of the *α*13 motion also explains why it was not observed in our atomistic simulation, even though an enhanced sampling scheme was employed. The motion of the helix *α*13 has clear implications for the dimerization process. A recent study has suggested that, besides the LG domain interface, the dimerization of hGBP1 also involves an interface between the two helices *α*13 [30]. The motion of *α*13 observed in our simulations leads to an increased proximity of the two helices in the dimer (Fig 7). While the distance between the two *α*13 helices that we monitored already agrees with experimental findings, despite only one of the two helices having moved, it is likely that both *α*13 helices adopt the 90° rotated position in the dimer. Moreover, it is not implausible that dimers are preferentially formed by monomers which already have both *α*13 helices rotated, as the *α*13 motion is more likely to occur in the monomer than in the dimer. This is due to a newly formed salt bridge between *α*4’ and *α*13 in the dimer, which increases the energy barrier for the motion of *α*13. The rotated *α*13 helices would form a protein–protein interface, in addition to the LG domain interface involving the two guanine caps, which would further stabilize the hGBP1 dimer, as had already been suggested by Herrmann and co-workers [27, 30]. However, based on their DEER and FRET findings they proposed that *α*12 and *α*13 detach from the LG domain in order to allow for the two *α*13 helices coming into contact with each other [30]. Our simulation results demonstrate that such a detachment is not necessary to explain their experimental observations. In fact, such detachment would hinder the membrane binding of hGBP1 as our initial simulations of membrane-bound hGBP1 indicate (Fig 8E, unpublished results).

In summary, to understand the conformational flexibility of hGBP1 and its implication for the dimerization process, we used multi-resolution MD simulations in explicit solvent combined with PCA and MSM analysis. Our results indicate a hinge at the middle of the M and E domains leading to large-scale, dynamin-like motions, and highly flexible loops in the LG domain that open and close the nucleotide binding pocket without a nucleotide being present. We have further observed, for the first time to our knowledge, the change in orientation of helix *α*13 on a time scale of hundreds of *µ*s with direct implications for the dimerization of hGBP1. One possible scenario is that monomers that already have the helix *α*13 oriented away from helix *α*12 form a dimer where the two helices *α*13 are close enough to form an interface. Thus, the hGBP1 dimer, with interfaces between the LG domains and the helices *α*13, would be able to insert into a lipid membrane and, in combination with the motions of the M and E domains observed here, lead to the disruption of the membrane and so to the biological function of hGBP1.

## Methods

All software and web databases used in this work are listed in Table S1. The input needed by the various software and output files created are described in a README file in the Supporting Information.

### All-atom simulations

For all atomistic simulations of this study the Amber99SB^*^-ILDNP force field [37, 38] combined with the explicit water model TIP3P [39] were employed. Electrostatic interactions were treated with the particle-mesh Ewald method [40, 41] in conjunction with periodic boundary conditions and a real-space cutoff of 12 Å. The Lennard-Jones interactions were cut at 12 Å. A leapfrog stochastic dynamics integrator was used for the integration of equations of motion. The LINCS algorithm [42] was used to constrain all bond lengths and the hydrogen atoms were treated as virtual interaction sites, permitting an integration time step of 4 fs while maintaining energy conservation [43].

The crystal structure of the hGBP1 monomer in ligand-free form with PDB ID 1DG3 was used as starting conformation [16]. Missing amino acids from loops in the crystal structure were added with the software ModLoop [44, 45]. The final conformation was placed in a dodecahedral box with 12 Å between the protein and the box, solvated with 108,406 water molecules and 7 Na^+^ ions were added for charge neutrality, resulting in a system with a total number of 335,553 atoms. This particular simulation box was large enough to allow free translation and rotation of the hGBP1 protein without interacting with its periodic images that would otherwise result in simulation artifacts. After energy minimization and equilibration of the system following the same procedure as described in the next paragraph, a Hamiltonian replica exchange MD simulation [46] with 30 replicas was performed. The energy function of hGBP1 including hGBP1–water interactions was modified in each replica by applying biasing factors of 310 K/*T* with the 30 temperatures *T* exponentially distributed between 310 and 450 K. This implies one unbiased replica, the so-called target replica at 310 K. The average exchange probability between the replicas was ∼30%. Each replica simulation was 400 ns long, leading to a total of 12 *µ*s for the 30 replicas. The H-REMD simulation was realized with Gromacs 4.5.5 [47] in combination with the PLUMED plugin (version 2.1) [48].

The isolated E domain (residues 482–591) and the E domain plus helix *α*11 of the M domain (residues 456–591) were both simulated at the all-atom level for 500 ns using Gromacs 2016 [49, 50]. For the E domain two Cl^−^ ions and for *α*11 plus the E domain one Na^+^ ion were addded for neutrality of the systems. The energy of both systems was first minimized using a steepest descent algorithm, followed by equilibration of the systems to the desired temperature of 310 K and pressure of 1 atm for mimicking the physiological environment. First, a 0.1 ns *NV T* equilibration was performed in which the number of atoms (*N*), the box volume (*V*) and temperature (*T*) were kept constant, followed by a 1 ns *NpT* equilibration to adjust the pressure (*p*). During equilibration, the protein atoms were restrained with a force constant of 10 kJ mol^−1^ Å^−2^ allowing the water molecules to relax around the solute. Finally, the 500 ns MD production runs in the *NpT* ensemble were performed. As no restraints were applied to the protein during the simulations of the E domain (with and without *α*11), a large cubic box with an edge length of 180 Å was created, allowing free rotation and translation of the 120-Å long helix *α*12 in all directions. The resulting system sizes involved about 573,000 atoms. The velocity rescaling thermostat was employed to regulate the temperature in the *NV T* simulations, while the Nosé-Hoover thermostat [51, 52] and the isotropic Parrinello-Rahman barostat [53] were used for the *NpT* simulations.

### Coarse-grained simulations

The coarse-grained simulations were performed with the Martini force field 2.2 and the Martini explicit water [21] as implemented in Gromacs 4.5.5 [47]. As initial conformation for the hGBP1 monomer we used the crystal structure with PDB code 1DG3 [16], which was inserted in a rectangular box with edge lengths of 85, 90 and 170 Å. The box was then solvated with 17,118 Martini water molecules and 7 Na^+^ ions resulting in a system with 18,438 particles. In order to avoid overall translation and rotation of the protein in the box, the stable secondary structure elements of the LG domain, which were identified based on the atomistic H-REMD simulation results, were restrained to their original positions. After an initial energy minimization and short equilibration, five MD simulations of the system at a temperature of 310 K were performed for different lengths, ranging from 63 *µ*s to 200 *µ*s (see Table 2).

Similar Martini simulations were conducted for the hGBP1 dimer. The dimer was built by superimposing the LG domain of the apo monomer (PDB ID 1DG3) to one of the LG domains (residues M1–L309) of the LG dimer in complex with GDP, which was resolved by crystallography (PDB 2B92) [12]. In order to avoid clashes between atoms, we replaced the helix involving residues T133–F175 in the dimer with the same helix from the monomer (Fig S7B). For completing the full dimer model, we combined the LG domain dimer up to V288 with the monomer conformation starting from N289. The resulting dimer model was first subjected to energy minimization at atomistic resolution and in explicit water, and then converted to the Martini model. The coarse-grained hGBP1 dimer was inserted in a rectangular box with edge lengths of 284, 111 and 108 Å filled with Martini water and 14 Na^+^ ions, amounting to a final system with 32,836 particles. Similarly to the Martini monomer simulation, the stable parts of the LG domain were restrained for both proteins in order to avoid the dimer to rotate and translate, and also to preserve the conformation of the LG domains. In addition, also the G2-SW1 and GC loops were restrained to keep the LG domains in its dimer-specific conformation as present in PDB 2B92 [12]. As listed in Table 2, for the dimer system we performed 5 simulations at 310 K (between 23 *µ*s and 150 *µ*s of length) and one simulation at 320 K (270 *µ*s).

### Analysis

To create pictures of the 3D protein structures, the Visual Molecular Dynamics (VMD) software [55] and PyMol [56] were used. If not stated otherwise, for the analysis of the H-REMD simulation the data collected by the target replica was used. To quantify the stability and flexibility of hGBP1 during the atomistic MD simulations, the root mean square fluctuations (RMSF) of the C_*α*_ around their average positions was calculated. The RMSF calculated for different time intervals of the H-REMD simulation was used to demonstrate that this simulation had converged within 400 ns per replica (Fig S8). To determine the time-resolved secondary structure of the E domain the DSSP algorithm (Define Secondary Structure of Proteins) [57] was employed. Clustering analyses were performed to obtain the most populated conformations of the E domain and loops of the LG domain using the Daura algorithm [58] applied to all atoms of the structural element in question and cut-off values of 3.5 Å for the E domain and 2.5 Å for the loops of the LG domain. The details for the calculation of distance maps are given in the Results.

The main structural changes of the M and E domains sampled by the target replica of the H-REMD simulation were identified based on a principal component analysis (PCA) [59]. Given the existence of many different structural fluctuations in hGBP1, we found that applying the PCA to the entire protein is not the best way for separating the different large-scale motions of the protein from each other. Therefore, we only considered the Cartesian coordinates of the M domain and *α*12 during that analyis as the RMSF analysis had revealed that these regions exhibit the largest flexibility. Helix *α*13 of the E domain was not included as it was very stable throughout the H-REMD simulation. We projected the conformations from the H-REMD target replica onto the first two principal components (PCs), calculated two-dimensional histograms and then the 2D free energy surface along the two PCs.

For the analysis of the coarse-grained simulations of the hGBP1 monomer, we applied the Markov state model (MSM) approach using the PyEmma software [60]. First, the five trajectories were subjected to the time-lagged independent component analysis (TICA) [61], a method well suited for dimensionality reduction and recently applied with success in the field of MD simulations [62–65]. The variance of the first two time-lagged independent components (TICs) amounted to 27% of the total variance, and they described best the conformational change involving helix *α*13. The MSM was then built by clustering the trajectories projected onto the first two TICs using the uniform time clustering algorithm with 300 microstates. We estimated the implied time scales from the MSM for 10 different lag times, based on which we selected a lag time of 450 ns for caclculating the MSM of our system. For the identification of metastable Markov states we applied the fuzzy spectral clustering method PCCA+ [66, 67] and used transition path theory [68–70], which is implemented in the PyEmma software, to calculate the reactive fluxes yielding the mean first passage times between the states.

## Supporting information

**S1 Fig. Information on the eigenvalues and eigenvectors obtained from the PCA of the all-atom H-REMD simulation.** (A) Amplitude of the first ten eigenvalues obtained from principal component analysis (PCA) of the all-atom H-REMD simulation of the hGBP1 monomer. (B) Projection of the unbiased H-REMD replica onto the first five PCA eigenvectors. (PDF)

**S2 Fig. Dynamics of the isolated E domain of hGBP1 obtained from all-atom MD simulations.** (A) Representative conformations for clusters 3–5 with their occurrence (in %) obtained from the MD simulations of the isolated E domain (top) and of helix *α*11 plus the E domain (bottom). The cluster structures were aligned to the crystal structure of full-length hGBP1 (shown as transparent cartoon) using residues 482–484 for the alignment of the isolated E domain and *α*11 for alignment of *α*11 plus the E domain. (B) Evolution of the secondary structure shown for each residue as a function of time for the isolated E domain (top) and for helix *α*11 plus E domain (bottom). (PDF)

**S3 Fig. Dynamics of the hGBP1 monomer characterized by Martini MD simulations.** (A) Free energy surface plotted along the first two eigenvectors obtained from TICA. The free energy values correspond to the color scale on the right (in *k*_B_*T* with *k*_B_*T* as the Boltzmann constant and *T* = 310 K). (B) Fuzzy PCCA+ clustering of the microstates resulted in six macrostates. The microstates are projected onto the first two TICs and their membership to one of the six macrostates is identified by different colors. (PDF)

**S4 Fig. Implied time scales obtained from Markov state modeling.** (A) Convergences of the implied timescales derived from MSMs for different lag times. One step corresponds to 1.5 ns. (B) Relative implied time scales for a lag time of 450 ns. (PDF)

**S5 Fig. Contact maps for the interactions between** *α***13 and the LG domain.** (A) Distance map between the C_*α*_ atoms of the C-terminal region, including the helix *α*13, and the loop formed by residues R151–F174 during transition from the Markov state A to other states in the MSM. The residue index on the *x* and *y* axes is accompanied by the corresponding amino acid sequence on the respective opposite side of the plot. (B) C_*α*_-distance map between the C-terminal region as in (A) and relevant residues of the LG-domain calculated for conformations belonging to the Markov state 4. The sequences of the LG domain stretches that form strong contacts with the C-terminal region are displayed on the right, together with structural snapshots where the same residues are highlighted in red, while the helices *α*12 and *α*13 are shown on blue and cyan, respectively. (PDF)

**S6 Fig. Positions of** *α***3 and** *α***4’ in the hGBP1 monomer and LG domain dimer.** (A) Comparison of the position of helix *α*4’ from the hGBP1 monomer (PDB 1DG3, red) with that in the LG domain dimer (PDB 2B92, yellow). This is the major conformational difference between the LG domain monomer and dimer. (B) Comparison of the position of helix *α*3 in the LG domain dimer (PDB 2B92, yellow) with that from the hGBP1 monomer (PDB 1DG3, red). In order to avoid atom clashes when building the full-length hGBP1 dimer, the yellow helix had to be replaced with the red one. (PDF)

**S7 Fig. E217–K567 distance during selected coarse-grained simulations.** Distance between the side chains of E217 and K567 for representative Martini simulations of (black) the hGBP1 monomer at 310 K (obtained from the 63 *µ*s simulation), (red) and (blue) the hGBP1 dimer at 320 K (simulated for 270 *µ*s). In one of the monomers composing the dimer the helix *α*13 moves (blue) while it does not in the other monomer (red). This motion correlates to the presence of the salt bridge between E217 and K567. In the monomer this salt bridge is never formed (black) as these two residues are too far away from each other. In the dimer, helix *α*4’ adopts a slightly different position than in the monomer (see Fig S7), allowing a salt bridge being formed between E217 and K567. This salt bridge impedes the motion of *α*13 in the dimer. Only if the temperature is raised to 320 K, *α*13 starts moving in one of the monomers composing the dimer (blue) while it remains intact in the other one (red). At 50 *µ*s the salt bridge in the monomer with the flexible *α*13 is completely broken, which corresponds to the time when this helix has adopted the 90° rotated position (see Fig 7). It should be noted that the distances shown here are between the centres of the coarse-grained side-chain beads and are therefore larger than the atom-based distances usually reported for salt bridges. (PDF)

**S8 Fig. Convergence of the H-REMD simulations.** The fluctuations of hGBP1 during the first half of the H-REMD simulation (red), the second half (blue) and the full simulation (black) are almost identical, showing that no new conformations are sampled in the second half of the simulation. The results are shown as RMSF values calculated for the target replica (310 K, no energy bias). (PDF)

**S1 Table. Web resources–Software and databases.** (PDF)

**S1 README.** Description of the GROMACS and PyEMMA input / output files available via the Open Science Framework project account https://osf.io/a43z2/. (PDF)

## Acknowledgments

This project was funded by the Deutsche Forschungsgemeinschaft (DFG, German Research Foundation) – project number 267205415 – CRC 1208. The authors gratefully acknowledge the computing time granted through JARA-HPC (project JICS6A) on the supercomputer JURECA at Forschungszentrum Jülich.

## Author contributions

B.B. and B.S. designed the computational experiments, B.B. and J.L. conducted and analyzed the simulations. All authors discussed the results and wrote different parts of the manuscripts. B.S. reviewed the manuscript.

